# Cloning, Biochemical Characterization and Inhibition of Alanine racemase from *Streptococcus iniae*

**DOI:** 10.1101/611251

**Authors:** Murtala Muhammad, Yangyang Li, Siyu Gong, Yanmin Shi, Jiansong Ju, Baohua Zhao, Dong Liu

**Affiliations:** College of Life Science, Hebei Normal University, Shijiazhuang 050024, China

**Keywords:** *Streptococcus iniae*, Alanine racemase, Peptidoglycan, Homogentisic acid, Hydroquinone

## Abstract

*Streptococcus iniae* is a pathogenic and zoonotic bacteria that impacted high mortality to many fish species, as well as capable of causing serious disease to humans. Alanine racemase (Alr, EC 5.1.1.1) is a pyridoxal-5′-phosphate (PLP)-containing homodimeric enzyme that catalyzes the racemization of L-alanine and D-alanine. In this study, we purified alanine racemase from the pathogenic strain of *S. iniae*, determined its biochemical characteristics and inhibitors. The *alr* gene has an open reading frame (ORF) of 1107 bp, encoding a protein of 369 amino acids, which has a molecular mass of 40 kDa. The optimal enzyme activity occurred at 35°C and a pH of 9.5. The enzyme belongs to the PLP dependent enzymes family and is highly specific to L-alanine. *S.iniae* Alr can be inhibited by some metal ions, hydroxylamine and dithiothreitol (DTT). The kinetic parameters *K*_*m*_ and *V*_*max*_ of the enzyme were 33.11 mM, 2426 units/mg for L-alanine and 14.36 mM, 963.6 units/mg for D-alanine. Finally, the 50% inhibitory concentrations (IC_50_) values and antibiotic activity of two alanine racemase inhibitors, were determined and found to be effective against both gram positive and gram negative bacteria employed in this study. The important role of alanine racemase as a target of developing new antibiotics against *S. iniae* highlighted the usefulness of the enzyme for new antibiotics discovery.

## 1. Introduction

*Streptococcus iniae* (*S. iniae*) is a gram-positive and most commonly reported fish streptococcal pathogen responsible for high economic loses of aquaculture industries around the world. The zoonotic bacteria was also reported to cause bacteremia, cellulitis, meningitis, and osteomyelitis in human (Guo et al., 2018; Tavares et al., 2018). Vaccines and antibiotics were currently employed for minimizing the impact of the disease, however, recent studies revealed that the bacteria has so far developed resistance against many potential antibiotics (Tavares et al., 2018; Muhammad et al., 2019) as such, additional efforts for developing more effective vaccines and antibiotics are necessary steps for circumventing the threat of its infection (Saavedra et al., 2004). One promising target of new antibiotics discovery is alanine racemase.

Alanine racemase (Alr; E.C. 5.1.1.1) is an enzyme that catalyzes the interconversion of l-alanine and D-alanine using a pyridoxal 5-phosphate (PLP) as a cofactor (Tassoni et al., 2017). d-alanine was used for the synthesis of peptidoglycan of the bacterial cell wall. it is directly involved in cross-linking of adjacent peptidoglycan strands and also present in lipoteichoic acids of Gram-positive bacteria (Liu et al., 2018; Ray et al., 2018). There are two isoforms (non-homologous) of alanine racemase genes (*alr* and *dadX*). The *alr* gene which is constitutively expressed is an essential enzyme for cell wall synthesis while the expression of *dadX* is induced in the presence of high concentrations of L- or D-alanine. *DadX* is basically required for l-alanine catabolism, forming a substrate for d-alanine dehydrogenase (*dadA*) (Duque et al., 2017). The bacterial cell wall is indispensable for the survival and viability of bacteria (Liu et al., 2019) and has always been an interesting target for many antibiotics and antimicrobial agents (Anthony et al., 2011). Alanine racemase is ubiquitous among bacteria and rare in eukaryotes but absent in humans (Kawakami et al., 2018), hence it emerges as an attractive and potential therapeutic target for the antimicrobial drug (Wang et al., 2017).

Numerous inhibitors were identified as able to affect the activity of alanine racemase (Kim et al., 2003a; Kim et al., 2003b). Many of the inhibitors were structural analogs of alanine: they interact with the enzyme-bound PLP, covalently bound to some eukaryotic PLP-dependent enzymes and lead to cellular toxicity (Toney, 2005). PLP-related off-target effects could be overcome by using enzyme inhibitors that are not substrate analogs. Structure-based approach and molecular modeling have been employed to discover novel alanine racemase inhibitors which are devoid of affinity for the PLP and hence off-target effects (Lee et al., 2013; Azam and Jayaram, 2018).

In this study, we identified and purified the alanine racemase from *S. iniae* HNM-1 strain that was previously isolated from an infected Chinese sturgeon (*Acipenser sinensis*) after massive mortality as a result of its infection (Muhammad et al.,2019). We characterized its enzymatic properties, substrate specificity and kinetic parameters. We also explore the potentiality of the enzyme as an attractive antimicrobial target against *S. iniae*. We determined the 50% inhibition concentrations (IC_50_) of two alanine racemase inhibitors and their antimicrobial susceptibility against six opportunistic pathogens including *S. iniae*, in quest of providing the possible solutions against antibiotics resistance and bacterial infections.

## 2. Results

### 2.1. Identification of *S. iniae* alanine racemase gene

According to the genomic sequence of *S. iniae*, the bacteria have a single putative alanine racemase (*alr*) gene. The *alr* gene has an open reading frame of 1107bp that encodes a 369 amino acids protein (SiAlr) with a molecular mass of 39.82 kDa. The nucleotide sequence of *alr* has been submitted to GenBank under accession number MK620909.

The deduced amino acid sequence has 76%, 67%, 63% and 47% similarities with alanine racemase of *Streptococcus pyogenes, Streptococcus agalactiae, Streptococcus pneumoniae*, and *Enterococcus faecalis*, respectively. Phylogenetic analysis of alanine racemase from different bacteria revealed an evolutionary relationship among them. The phylogenetic tree consists of two distinct clades. The enzyme is clustered with other Streptococci species, such as *S. pyogenes, S. agalactiae, S. pneumoniae* and *E. faecalis*. These sequences, from gram-positive bacteria, were classified into one group. The sequences from gram-negative bacteria, such as *Pseudomonas aeruginosa, Aeromonas hydrophila* and *Corynebacterium glutamicum* were classified into another (Fig. 1). The evidence indicated that these alanine racemases evolved independently from a common ancestor and formed two isolated genes.

**Fig. 1.**
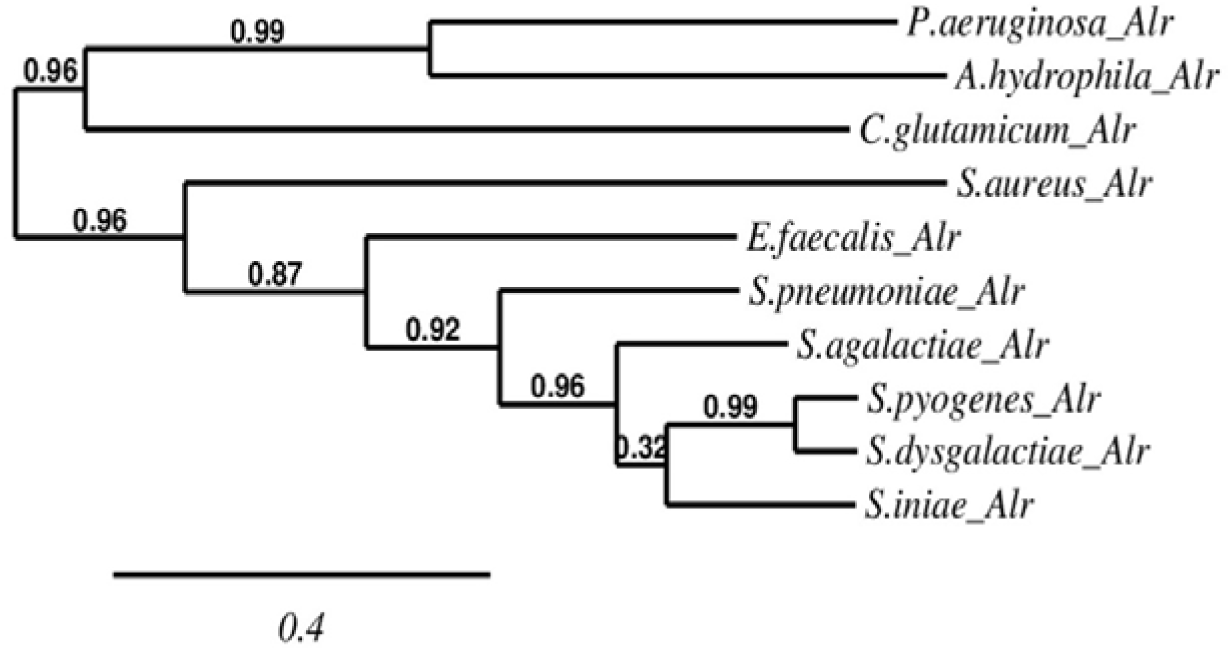
Phylogenetic relationships of SiAlr and sequences from 10 other species. The tree was constructed using Neighbor end-joining. Maximum likelihood tree based on complete coding sequences deposited in GenBank. The evolutionary distances were computed using the p-distance method while the scale bar indicates 0.4 amino acid substitutions per site.

Multiple sequence alignment of SiAlr with sequences of other 10 species suggested that some regions are absolutely conserved in SiAlr, which includes PLP binding motif near the N-terminus (AVVKANAYGHG) and the two catalytic amino acid residues of the active center (Lys 40 and Tyr 274). The eight residues making up the entryway to the active site (inner layer: Ala 174, Tyr 273, Tyr 282 and Tyr 366; middle layer: Asp 166, Arg 298, Arg 318 and Ile 364) (Fig. 2).

**Fig. 2.**
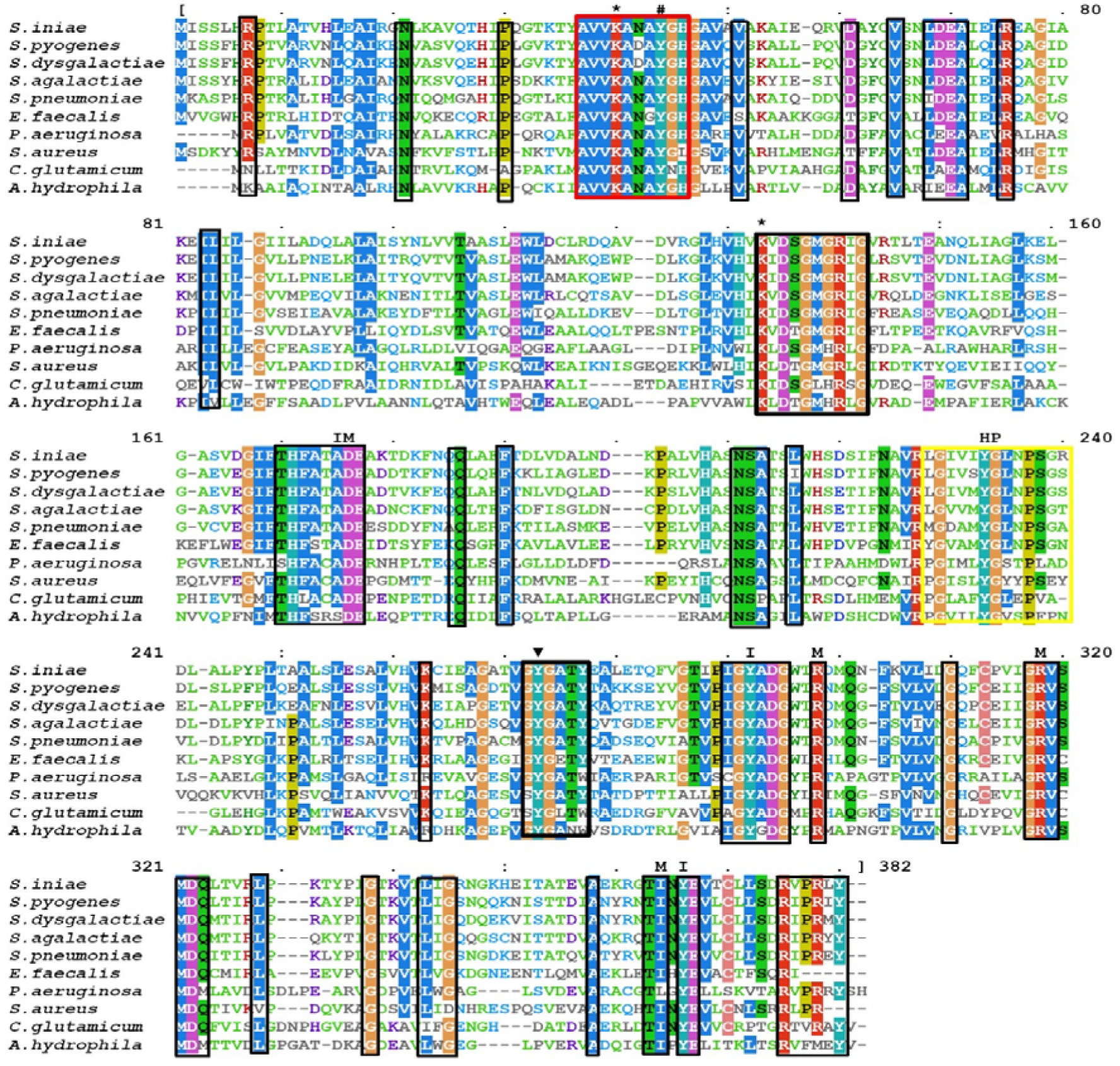
Structure-based sequence alignment of alanine racemases sequences. The amino acid sequence of Alr from *S. iniae* was aligned with alanine racemases sequences of *S. uberis* CAIM 1894, *S. agalactiae* 2603V/R, *E. feacalis* D32, *B. anthracis* H9401, *S. aureus* ABFQT, *C. glutamicum* ATCC 13032, *A. hydrophyla* ATCC 7966, *S. pneumoniae* MDRSPN001, and *E.coli* CVM N33720P. The red box enclosed the conserved PLP-binding sites; Lys40 (*), Tyr44 (#). The catalytic Tyr residue was indicated by (+). Strictly conserved residues were enclosed in the black boxes, while the hydrophobic patch (HP) in the yellow box. Residues of the active site entryway are marked with either I (inner layer) or M (middle layer). Highly conserved residues were indicated by the box and strictly conserved with (*).

### 2.2. Expression and Purification of SiAlr

The SiAlr protein was expressed in *E.coli* (DE3) incubated overnight at 16 °C. The was purified to homogeneity using Ni-agarose affinity chromatography. The protein has a relative molecular mass of 39.82 kDa as estimated by SDS-PAGE, which was similar to the calculated molecular mass. Calculated relative molecular mass and western blotting analysis using the anti-poly-His antibody confirmed that 39.82 kDa protein is SiAlr (Fig 3).

**Fig. 3.**
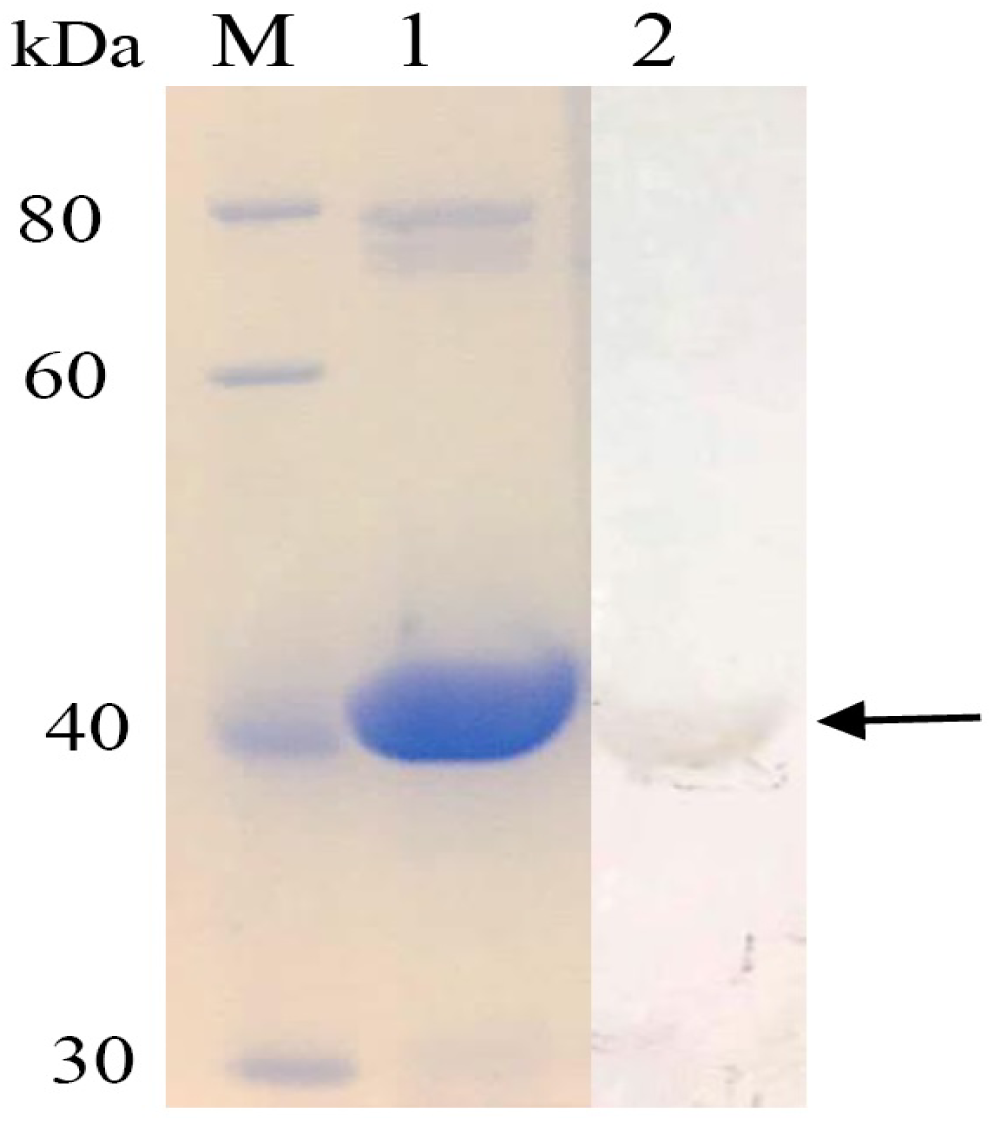
Purification of *S. iniae* alanine racemase. The enzyme was purified using Nickel ion affinity chromatography, analyzed by SDS-PAGE and western blotting. M: molecular weight standards; Lane 1: Protein marker, Lane 2; 40 kDa SiAlr. Lane 2: Western blotting analysis of the purified protein.

### 2.3. Characterization of the enzyme

The optimal temperature of SiAlr was approximately 35 °C. The enzyme was found to be very stable at the temperature of 30 and 35 °C, with more than 50% residual activity. The optimal pH of SiAlr was approximately 9.5 at 35°C. The enzyme was found to be very stable with more than 50 % residual activity after incubation for 2 hours at a pH of 8.5 to 9.5 (Fig 4). Thus demonstrating that SiAlr is a basophilic enzyme.

**Fig. 4.**
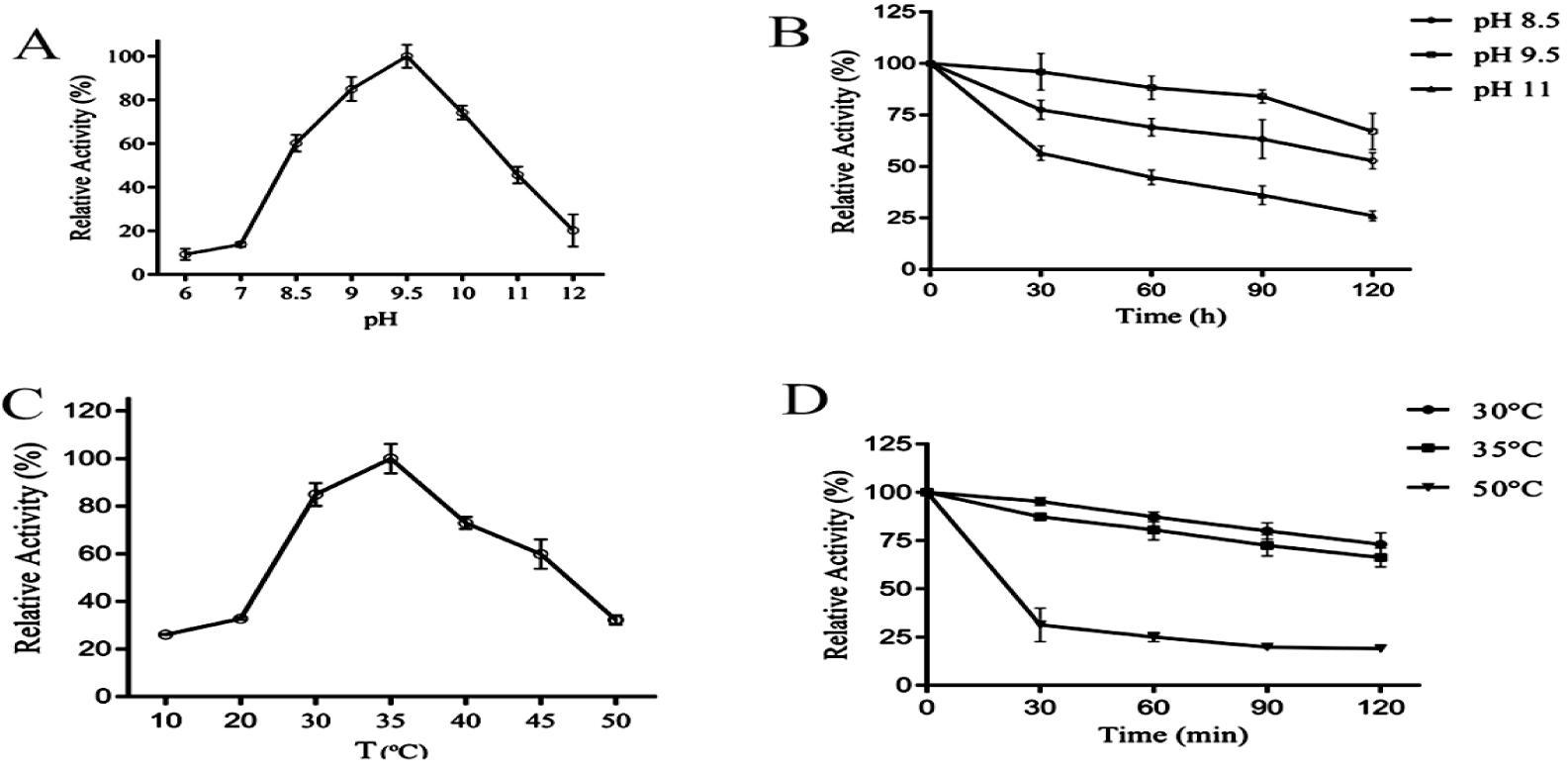
Effect of pH and Temperature on the activity of SiAlr. (A) Optimal pH; (B) pH stability; (C) Optimal temperature; (D) Thermal stability.

Various chemicals and metal ions were reported to inhibit the activity of alanine racemases. The results revealed that the enzyme activity was inhibited by most of the metal ions, but markedly inhibited by Ni^2+^, Co^2+^ Zn^2+^ and Fe^2+^ (Fig 5).

**Fig. 5.**
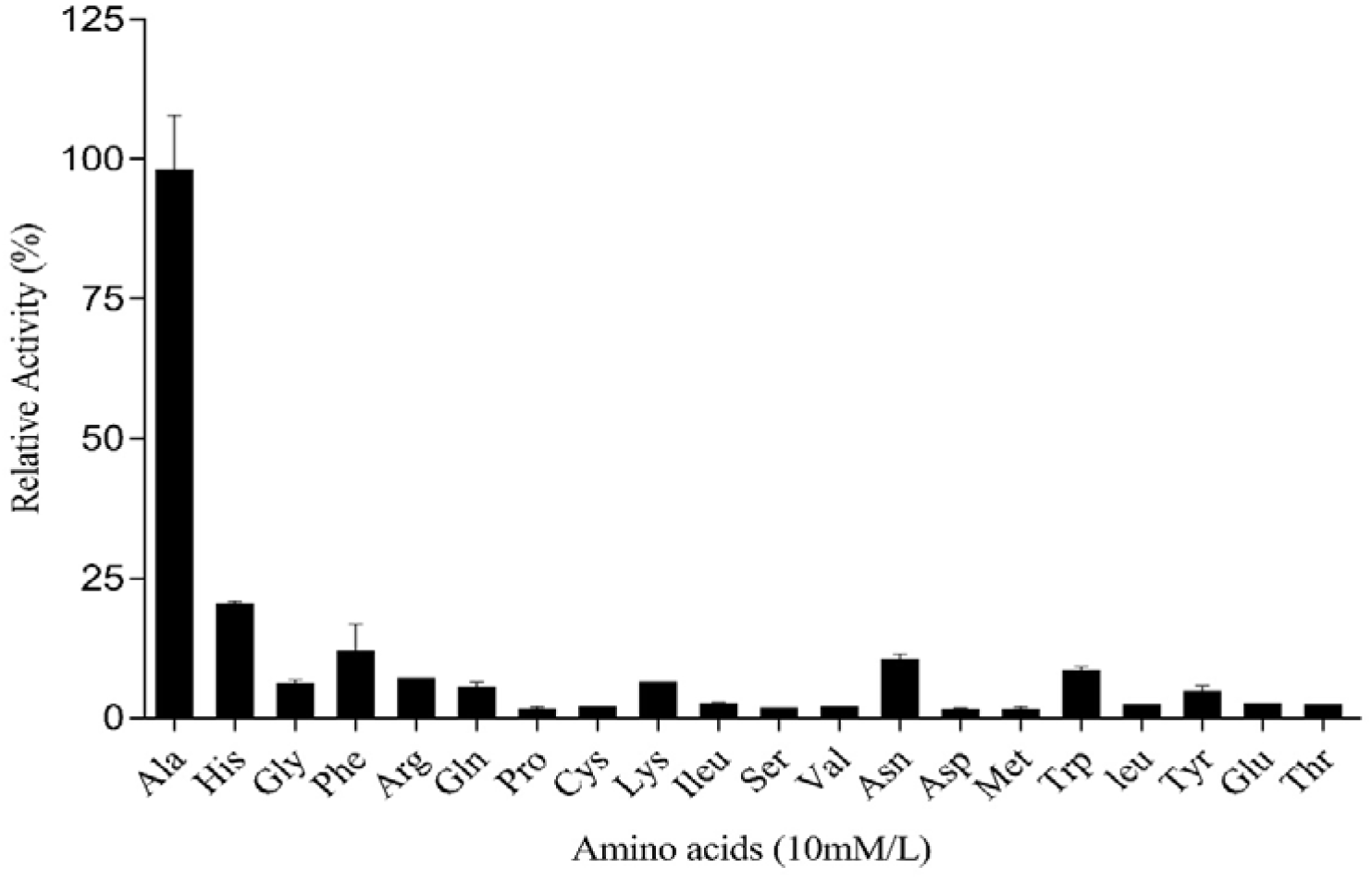
Effect of metals on SiAlr activity. The metal ions were at a concentration of 10mM/l. The data were presented as mean ± SD from 3 independent determinations.

### 2.4. Effect of reducing agents on the activity of SiAlr

Many inhibitors of alanine racemase have been discovered (Wang et al., 2017). The enzyme almost completely lost its activity after treatment with 1 and 10 mM hydroxylamine. Addition of 0.1mM hydroxylamine reduced the activity of the enzyme by 80%. Treatment of SiAlr with 1 mM/L of DTT resulted in a 70% loss of its activity and completely inhibited at the concentration of 3 mM (Table 2).

**Table 1.** Strains and plasmids used in this study.

| Strains/Plasmids | Description | Source |
| --- | --- | --- |
| **Strains** |  |  |
| *S. iniae* HNM-1 | isolated from infected *Acipenser sinensis* | This study |
| *E. coli* DH5α | used for cloning and propagation of plasmids | Novagen |
| *E. coli* BL21(DE3) | used for protein expression | Invitrogen |
| *Salmonella* |  | This study |
| *Staphylococcus aureus* |  | This study |
| *Acinetobacter baumannii* |  | This study |
| *Pseudomonas aeruginosa* |  | This study |
| **Plasmids** |  |  |
| pMD19-T | carries ampR gene; used for cloning PCR product with A at 3' ends | Takara |
| pET 22b (+) | carries ampR gene; used for expressing *S. iniae* Alanine racemase | Novagen |

**Table 2:** Effect of Hydroxylamine, DTT and PLP on SiAlr Activity.

| Chemical | Concentration, mM | Relative activity, % |
| --- | --- | --- |
| None |  | 100(0.7) |
| Hydroxylamine | 0.1 | 21(1.2) |
|  | 1 | 11(0.8) |
|  | 10 | 9(1.4) |
| DTT | 1 | 27(3.1) |
|  | 3 | 2(0.8) |
| PLP | 0.01 | 56(2.4) |
|  | 0.04 | 83(1.5) |
|  | 0.06 | 96(2.7) |

The data were presented as mean (SD) from 3 independent enzyme reactions.

We examined the role of PLP in the activity of SiAlr by resolving the enzyme to Apo-enzyme by Hydroxylamine treatment. The Apo-enzyme almost completely lost its activity after treatment with 10mM hydroxylamine. Addition of 0.01, 0.04 and 0.06mM PLP make the enzyme regained up to 56 %, 83% and 96% of its activity, respectively. The result indicated that SiAlr is a PLP-dependent enzyme that requiring more than 0.01 mM PLP to maintain its activity (Table 2).

### 2.5. Substrate specificity

Alanine racemase is a highly conserved bacterial enzyme and known to be very specific to its substrate (Patrick et al., 2002). As shown in Fig 6, the enzyme was highly specific to L-alanine and showed week activity with L-phenylalanine (11%), L-Histidine (20%), and L-Asparagine (10%). This result indicates that SiAlr has strict substrate specificity.

**Fig. 6.**
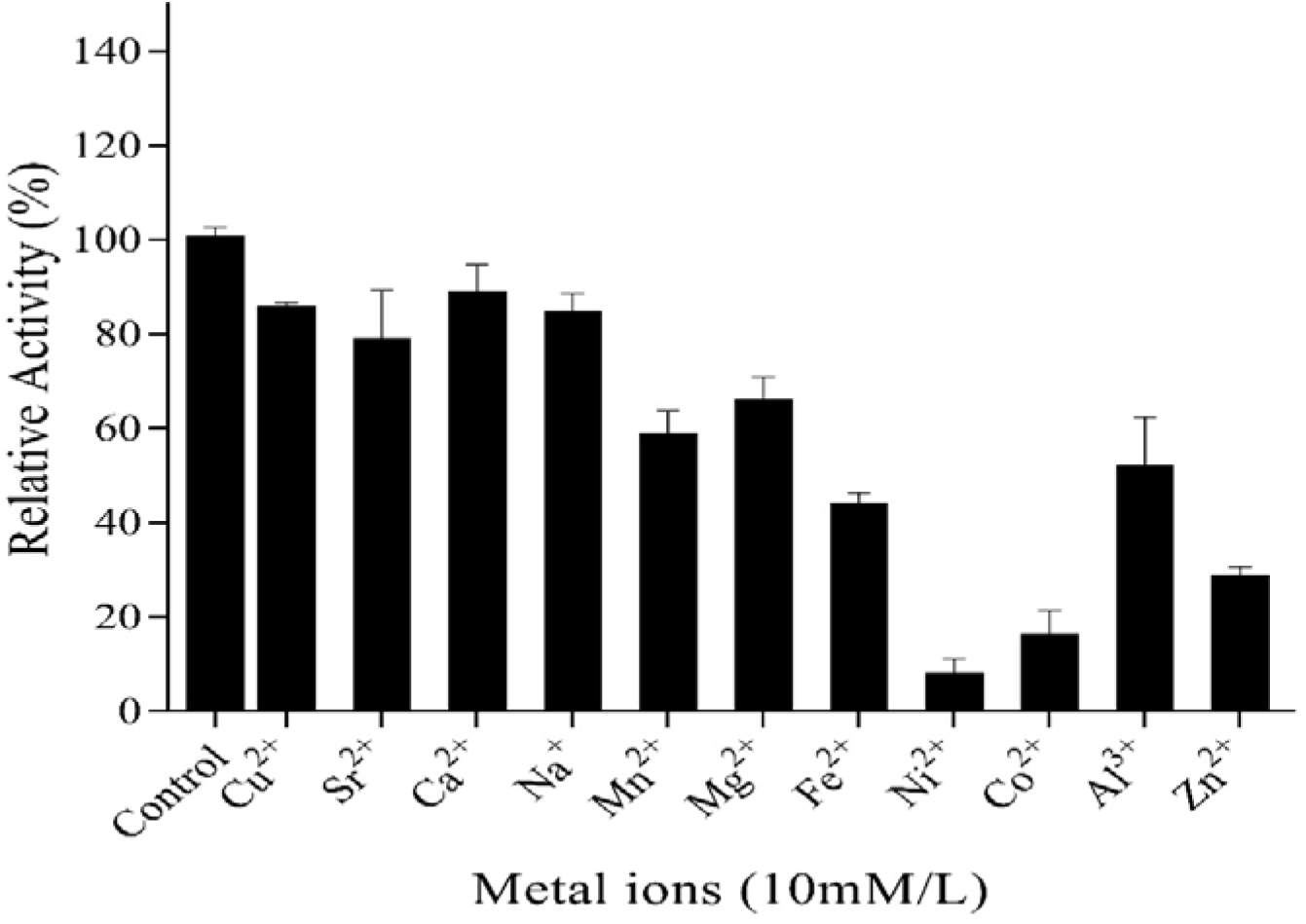
The substrate specificity of SiAlr. The relative activity of SiAlr for various L-Amino acids was determined at optimum pH and temperature. The data were presented as mean ± SD from 3 independent enzyme assays.

### 2.6. Kinetic parameters determination

Kinetic parameters of SiAlr were determined using HPLC. The substrate affinity constant (*K*_*m*_) for L-alanine was 33.11mM with a maximal velocity (*V*_*max*_) of 2,426 units/mg, while the D-alanine *K*_*m*_ value was 14.36 mM with a *V*_*max*_ of 963.6 units/mg. The *V*_*max*_ of L-alanine was more than twice than that of its enantiomer. These indicated that the enzyme has a greater binding affinity for L- alanine than for D-alanine, and the conversion of L- to D-alanine was more rapid than the reverse conversion. The equilibrium constant (*K*_*eq*_ (L/D)) was 1.09, which is consistent with the reported theoretical equilibrium constant (*K*_*eq*_ = 1) for alanine racemase (Liu et al., 2015) (Table 3).

**Table 3.** Kinetic parameters of SiAlr. The parameters were determined using HPLC and analyzed with Graph prism 6.0. The data were presented as mean ± SD from 3 independent enzyme assays.

| Enzyme | L to D-alanine | | D to L-alanine | | $K_{eq}(L/D)$ |
| --- | --- | --- | --- | --- | --- |
| | $K_m$ (mM) | $V_{max}$ (units/mg) | $K_m$ (mM) | $V_{max}$ (units/mg) | |
| SiAlr | 33.11 | 2426 | 14.36 | 963.6 | 1.09 |

**Table 4.** The results of antimicrobial activity of homogentisic acid and hydroquinone inhibitors against numerous isolates of gram-positive and gram-negative bacteria.

| Organism | <sup>a</sup>MIC (µg/ml) |  |
| --- | --- | --- |
|  | Hydroquinone | Homogentisic Acid |
| *S. iniae* HNM-1 | 25(2.3) | 200(5.6) |
| *Escherichia coli* DH5α | 130(7.9) | 210(8.4) |
| *Salmonella* | 150(8.7) | 180(11.4) |
| *Staphylococcus aureus* | 210(13.7) | 250(14.1) |
| *Acinetobacter baumannii* | 180(11.5) | 210(12.3) |
| *Pseudomonas aeruginosa* | 0 | 0 |

<sup>a</sup>MIC, average values with standard deviations.

### 2.7. IC_50_ determination

In our previous study, we found that homogentisic acid and hydroquinone are two alanine racemase inhibitors with minimal cytotoxicity against mammalian cells and can be utilized as potential agents of antibiotics (Wang et al., 2017). In this study we investigated the inhibitory effects of homogentisic acid and hydroquinone on SiAlr, DMSO was used as the blank control and DCS, a known alanine racemase inhibitor, as the positive control. The results showed that the IC_50_ values of hydroquinone and homogentisic acid were 11.39µM and 12.27µM. The IC_50_ values of hydroquinone and homogentisic acid were 3 and 3.3 times higher than that of DCS (Fig 7).

**Fig. 7.**
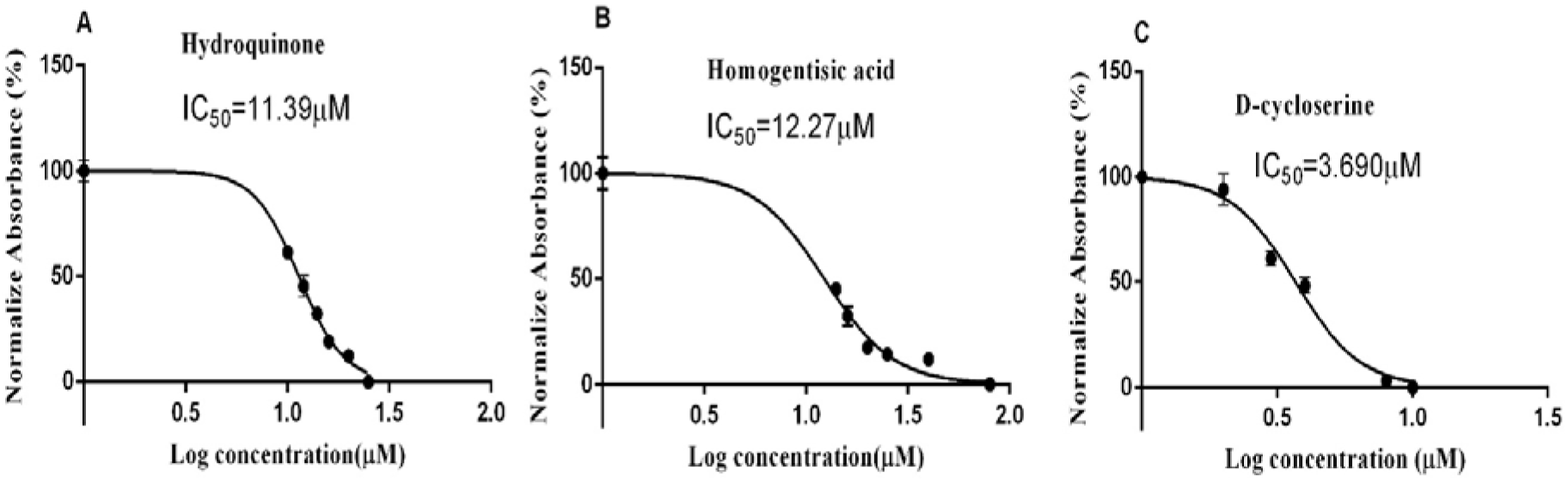
IC_50_ of the *S. iniae* alanine racemase inhibitor. A: IC_50_ of Hydroquinone was 11.39µM; B: IC_50_ of homogentisic acid was 12.27µM; C: IC_50_ of D-cycloserine was 3.69µM. The data shown are the means from three independent experiments.

### 2.8. Antimicrobial activity of alanine racemase inhibitors

The minimum inhibitory concentration (MIC) was conducted to determine the antimicrobial activity of two alanine racemase inhibitors against *S. iniae* HNM-1 and several conditional pathogenic bacteria. The results showed that hydroquinone and homogentisic acid have broad-spectrum antibiotic activities against both gram-positive and gram-negative bacteria. Hydroquinone showed good antibiotic activity against *S. iniae* HNM-1 with MIC value of 25 µg/ml, however, showed moderate antibiotic activity against other strains with MIC value of 130-210 µg/ml. Homogentisic acid demonstrated moderate antibiotic activity against bacteria tested with MIC value of 180-250µg/ml. Interesting, hydroquinone and homogentisic acid had no antibiotic activity against *Pseudomonas aeruginosa*.

## 3. Discussion

*S. iniae* is a gram-positive and one of the leading fish pathogens responsible for causing morbidity and mortality of more than 30 fish species (Aruety et al., 2016). *S. iniae* HNM-1 strain was isolated from an infected Chinese sturgeon (*Acipenser Sinensis*) after a disease outbreak that causes high morbidity and mortality (Muhammad et al., 2019). The classification of *S. iniae* HNM-1 was confirmed by molecular analysis of *16s rRNA* gene sequence. The sequence was deposited at the NCBI Genbank database under accession number KY781829. According to the genomic sequence of *S. iniae* 89353 strain (NCBI accession number CP017952.1), the bacteria have a single putative alanine racemase (*alr*) gene. As reported earlier, most of the Gram-positive bacteria, including *Lactobacillus plantarum* (Palumbo et al., 2004), *Bacillus anthracis* (Couñago et al., 2009), *Mycobacterium tuberculosis* (Nakatani et al., 2017) and *Mycobacterium smegmatis* (Chacon et al., 2002), appear to possess only one alanine racemase gene.

The optimal pH and temperature of SiAlr were 9.5 and 35°C respectively, which were similar to alanine racemase from *Aeromonas hydrophila* (Liu et al., 2015) and *Bacillus pseudofirmus* OF4 (Ju et al., 2009). Nearly all characterized alanine racemases have optimal pH more than 8 including enzymes from acidophile, *Acidiphilium organovorum* and *Acetobacter aceti* (Seow et al., 1998; Francois and Kappock, 2007). SiAlr is a mesophilic enzyme, stable at a temperature of 0°C to 40°C. Thermal stability of an enzyme is correlated with the host bacteria physiology and environment, thermophilic bacteria Alr are more stable than mesophilic and psychrotroph bacteria (Soda and Tanizawa, 1990; Yokoigawa et al., 1993). The effects of metal ions and other reagents on enzymes are diverse. Studies of alanine racemase from *A. hydrophila* have revealed that divalent cations, such as Ca^2+^ and Mg^2+^ enhanced racemization of alanine racemase (Liu et al., 2015). SiAlr was characterized as PLP-dependent racemase and showed high substrate specificity to alanine which is similar to most of the characterized Alanine racemases (Kawakami et al., 2018).

Many studies have focused on alanine racemase to develop antibacterial drugs for multiple bacterial species (Scaletti et al., 2012; Shrestha et al., 2017). Unlike D-cycloserine which is a cyclic analog of alanine and exerted its inhibitory effect through interaction with the enzyme-bound PLP cofactor (Batson et al., 2017), both homogentisic acid and hydroquinone are not structural analogs of Alr as such they are not interfering with other PLP dependent enzymes (Wang et al., 2017). According to the results of antimicrobial activity assay, the two inhibitors are capable of inhibiting both gram-positive and gram-negative bacteria with varying efficacies except *Pseudomonas aeruginosa*. The reason for two compounds showed no antimicrobial activity against *P. aeruginosa* may be that homogentisic acid is a normal product of *P. aeruginosa* and *P. aeruginosa* contained hydroquinone oxidase that oxidized hydroquinone (Higashi, 1958; Hunter and Newman, 2010).

Several alanine racemases have been identified and characterized form the *Streptococcus* species. Alanine racemase from *Streptococcus faecalis* NCIB 6459 with the molecular weight of 42kDa was the first one that was purified and characterized (Badet and Walsh, 1985). Strych et al isolated and characterized the alanine racemase gene from *Streptococcus pneumoniae*. They obtained preliminary crystals of *S. pneumoniae* Alr, and intend to incorporate the enzyme into the structural-based drug design program (Strych et al., 2007). Im et al solved the structure of *S. pneumoniae* Alr and identified three regions on the enzyme that could be targeted for structure-based drug design (Im et al., 2011). Qiu et al. first provided the first evidence that D-Ala metabolism is essential for planktonic growth and biofilm formation of *Streptococcus mutans*. It would be possible to take Alr of *S. mutans* as an antibacterial target to screen and optimize the safety and effective specificity of agents (Qiu et al., 2016). Wei Y. et al. confirmed that *alr* is an essential factor to maintain the growth and cell wall integrity of *S. mutans* (Wei et al., 2016). A serial of in vivo and in vitro experiments demonstrate that Alr is essential for the cariogenicity of *S. mutans*. Alr might represent a promising drug target to control the prevalence of cariogenic *S. mutans* in a multi-species microbial consortium and be a potential target for the prevention and treatment of caries (Liu et al., 2018). Therefore, Alr is regarded as a drug target for further investigation to develop effective drugs against *S. iniae* and a subject of mutational studies for the development of mutants with enhance activity that can be utilized for industrial purposed, since d-alanine is also widely used for the production of infusion solutions (Nachbauer et al., 1984), food additive (Awasthy et al., 2012), and in the manufacturing of artificial fibers (Teulé et al., 2009).

### 3.1. Conclusion

Purification and characterization of Alr from both gram-positive and gram-negative bacteria is an essential step towards an in-depth understanding of enzyme divers features, design new broad-spectrum antibiotics and used for site-directed mutagenesis in order to improve the enzyme catalysis and stability. Hydroquinone and homogentisic acid are promising inhibitors of Alr that are capable of inhibiting the growth of both gram-positive and gram-negative bacteria. Future investigation will focus on finding the physiological role of Alr, exploring new novel antimicrobial agents against *S. iniae* and improving their efficacy by designing and analyzing their new derivatives that may have improved antimicrobial activity.

## 4. Materials and Methods

### 4.1. Strains, plasmids, and growth conditions

The characters of bacterial strains and plasmids used in this context were summarized in Table 1. *S. iniae* HNM-1 was cultured at 35°C in Tryptone soy yeast extract (TSYE) medium. *Escherichia coli* (*E. coli*) DH5α, *E. coli* BL21 strains, *Salmonella, Staphylococcus aureus, Acinetobacter baumannii* and *Pseudomonas aeruginosa* were cultured in Luria Bertani (LB) media at 37°C or 35°C. For the final concentration of antibiotic, 100-g/mL ampicillin (Amp) was used in this study.

### 4.2. Cloning of alanine racemase gene

Primers were design based on the *alr* gene sequence of *S. iniae* 89353 strain (NCBI accession number CP017952.1). The genomic DNA of *S. iniae* HNM-1was extracted and amplified using the primers: Alr-F-(5’-GCA<u>CCATGG</u>ATGATTTCAAGTTTG-3’) and Alr-R-(5’-TCA<u>CTCGAG</u>ATCCCGATAAAGC-3’), with *Nco*I and *Xho*I restriction sites, underlined respectively. PCR product was cloned in pMD19-T cloning vector to construct pMDalr and transformed to *E. coli* DH5α. The *alr* gene was digested with restriction endonucleases and cloned into expression vector pET-22b (+), forming recombinant plasmid pET22b-*alr*, and subsequently, the gene was sequenced and analyzed. The deduced amino acid sequence of the ORF was analyzed by the Blast software. Multiple amino acid sequence alignment and phylogenetic relationships among alanine racemase of *S. iniae* HNM-1 (SiAlr) and other bacteria were constructed with Clustal Omega.

### 4.3. Expression and purification of alanine racemase

The expression vector was transformed into *E.coli* BL21 (DE3) for protein expression, a single colony of the transformed *E.coli* was inoculated in 100ml LB medium which contained ampicillin (100µg/ml) and incubated at 35°C. Protein expression was induced when the OD_600_ reaches 0.6 by addition of IPTG at a final concentration of 1mM, and re-incubated overnight at 16°C or at 35°C for 5 hours. Cells were collected and resuspended in 20 ml binding buffer (50 mM NaH_2_PO_4_, pH 8.0, 300 mM NaCl and 10mM imidazole), lysed on ice by sonication for 40 minutes, and centrifuged at 8000g, 4 °C for 10 minutes. The supernatant was collected and purified using Nickel ion affinity chromatography (Qiagen) according to the manufacturer’s protocol. The protein was dialyzed against phosphate buffered saline (PBS, pH 7.4). Protein purity and concentration were determined by SDS-PAGE and BCA protein assay kit (Takara) respectively. Western blotting was conducted using a monoclonal antibody against the poly-Histidine tag attached to the Alr protein as described previously (Liu et al., 2015).

### 4.4. Enzyme assay

Alanine racemase racemization assay was conducted in two coupled enzyme reactions, using standard racemization mixture (Wang et al., 2017). The reaction was initiated by addition of suitable concentration of SiAlr in the final reaction volume of 200 µl, incubated at 35 °C for 10 minutes, and terminated by addition of 25 µl of 2M HCl and neutralized with 25 µl of 2M sodium hydroxide, the reaction mixture was centrifuged at 14000 rpm, 4 °C for 10 minutes. The amount of D-alanine was measured in the second reaction containing 200 mM Tris- HCl pH: 8.0, 0.2 mg/ml 4-aminoantipyrine, 0.2 mg of N-ethyl-N-(2-hydroxy-3-sulfopropyl)-3-methyl aniline, 1unit of HRP, 0.1 unit of D-Amino acid oxidase and incubated at 37 °C for 20 minutes, The absorbance was measured using a microplate reader at 550nm.

### 4.5. Effect of temperature and pH on enzyme activity and stability

Influence of temperature was determined according to standard enzyme assay by measuring initial rate of reaction at various temperatures (10°C to 50°C), while the effect of pH was determined by measuring initial rate of reaction in Britton-Robinson buffer (pH 2.0 to 12.0) at optimum temperature. The relative residual activity was calculated with the highest activity as 100%. Thermal and pH stability were respectively determined by incubating enzyme in a reaction mixture without substrate at 30°C, 35°C, 40°C, and in buffers with pH range from 8.5 to 11.0 at 4°C for 2 hours, the reaction was initiated by addition of the substrate, and incubated at 35°C for 10 minutes. The relative activity was calculated using 0 hours sample activity as 100%.

### 4.6. Substrate specificity of alanine racemase

Substrate specificity of SiAlr was determined according to standard racemization reaction mixture using 18 kinds of L-amino acid as substrates and incubated at the optimum temperature for 10 minutes.

### 4.7. Effect of metal ions, reducing agents and PLP on enzyme activity

Effect of some metal ions and chemical compounds on the activity of the enzyme were determined by incubating the enzyme with them in the reaction mixture for 30 min, afterward, added the substrate and determined the relative residual activity according to the standard protocol (Liu et al., 2015).

Different concentrations of hydroxylamine (0.1, 1, and 10mM) and the enzyme were added to the reaction mixture without the substrate, dialyzed in phosphate buffered Saline for 40 min and determined its activity without the addition of PLP. The effect of Dithiothreitol (DTT) on the activity of SiAlr was determined by incubating the enzyme in different concentrations of DTT (1 and 3 mM) for 30 minutes and measured the relative activity. To confirm SiAlr is PLP dependent enzyme, the purified Alr was treated with 10 mM hydroxylamine and dialyzed to obtain the apoenzyme. The apoenzyme was incubated in different concentrations of PLP (0.01, 0.04 and 0.06 mM) and measured the relative activity.

### 4.8. Kinetic parameters

Alanine racemase activity was determined by measuring the amount of both enantiomers of alanine by high-performance liquid chromatography (HPLC) using a fluorescence detector and followed the method described earlier (Hashimoto et al., 1992). The reaction mixture containing 10µm PLP, 200mM carbonate buffer pH 9.5, and various concentrations of either L or D forms of alanine, followed by enzyme addition and incubation at 35°C for 10 min. The reaction was terminated by the addition of 40µl of 2M HCL on ice for 2 min, neutralized with 40µl 2M NaOH, and centrifuged at 10000g, 4°C for 5 min. A 40µl aliquot of the reaction was derivatized by addition of 280µl of 0.4M boric acid pH 9.0, 0.1% N-tert-butyloxy-carbomyl-L cysteine (Sigma) and 0.1% O-phthaldehyde. One unit of the enzyme was defined as the amount of enzyme that catalyzed the formation of 1µmol of L or D-alanine from either enantiomer per minute. Graph Pad Prism 6.0 was used for results analysis.

### 4.9. Enzyme IC_50_ determination

Inhibitory effects of homogentisic acid and hydroquinone on the activity of alanine racemase were determined as described previously (Wang et al., 2017). Fivefold dilution series (in DMSO) was prepared for the compounds, and the solutions were added to the wells of a 96-well plate to yield the final inhibitory concentrations. Each concentration was tested in triplicate. The substrate was added after incubation for 30 min, and the fluorescence intensity was measured after the reaction. The negative control was prepared without adding chemicals to the control wells and the D-cycloserine (DCS) was used as the positive control. Percentage inhibition at each inhibitor concentration was calculated with respect to the negative control. GraphPad Prism 6.0 was used for the calculation of the concentration that causes 50% inhibition (IC_50_).

### 4.10. Antimicrobial susceptibility tests

Minimum inhibition concentrations (MIC) of hydroquinone and homogentisic acid against both Gram positive and Gram negative bacteria were determined by microdilution assay according to the guidelines of the Clinical and Laboratory Standards Institute, document M31-A3 (CLSI., 2008), as described previously (Dal Pozzo et al., 2011). An overnight culture was subculture to OD_600_ of 0.3, diluted tenfold, five times. Aliquots were spread on agar plates in triplicate to determine the number of colony-forming units (CFU)/ml. Compounds were diluted in DMSO at concentrations of 200, 100, 80, 40, 20, or 10 μg/ml. DMSO solvent was used as a negative control of growth inhibition and DMSO alone was used as the blank control. All tests were performed in triplicate. The inoculum of bacteria in culture medium (100 μl; 10^5^ CFU; OD_600_ = 0.3) was added to each well-containing compounds and incubated at 30°C for 48 hours. MIC values were determined as the lowest concentration at which no growth was observed upon visual inspection after incubating.

## Acknowledgment

This work was supported by the Natural Science Foundation of Hebei Province (C2013205103); the Outstanding Youth Foundation of Department of Education of Hebei Province (YQ2014026); the Research Fund of Hebei Normal University (L2016Z03); the State Key Laboratory of Pathogen and Biosecurity (Academy of Military Medical Science) (SKLPBS1529); and the Science and technology research project of Hebei Normal University (ZD2018070).

## Reference

Anthony, K.G., Strych, U., Yeung, K.R., Shoen, C.S., Perez, O., Krause, K.L., Cynamon, M.H., Aristoff, P.A. and Koski, R.A., 2011. New classes of alanine racemase inhibitors identified by high-throughput screening show antimicrobial activity against *Mycobacterium tuberculosis*. PLoS One 6, e20374.

Aruety, T., Brunner, T., Ronen, Z., Gross, A., Sowers, K. and Zilberg, D., 2016. Decreasing levels of the fish pathogen *Streptococcus iniae* following inoculation into the sludge digester of a zero-discharge recirculating aquaculture system (RAS). Aquaculture 450, 335–341.

Awasthy, D., Bharath, S., Subbulakshmi, V. and Sharma, U., 2012. Alanine racemase mutants of *Mycobacterium tuberculosis* require D-alanine for growth and are defective for survival in macrophages and mice. Microbiology 158, 319–327.

Azam, M.A. and Jayaram, U., 2018. Induced fit docking, free energy calculation and molecular dynamics studies on *Mycobacterium tuberculosis* alanine racemase inhibitor. Molecular Simulation 44, 424–432.

Badet, B. and Walsh, C., 1985. Purification of an alanine racemase from *Streptococcus faecalis* and analysis of its inactivation by (1-aminoethyl) phosphonic acid enantiomers. Biochemistry 24, 1333–1341.

Batson, S., de Chiara, C., Majce, V., Lloyd, A.J., Gobec, S., Rea, D., Fül↕p, V., Thoroughgood, C.W., Simmons, K.J. and Dowson, C.G., 2017. Inhibition of D-Ala: D-Ala ligase through a phosphorylated form of the antibiotic D-cycloserine. Nature communications 8, 1939.

Chacon, O., Feng, Z., Harris, N.B., Cáceres, N.E., Adams, L.G. and Barletta, R.G., 2002. *Mycobacterium smegmatis* D-alanine racemase mutants are not dependent on D-alanine for growth. Antimicrobial agents and chemotherapy 46, 47–54.

Clinical Laboratory Standards Institute. Antimicrobial disk and dilution susceptibility tests for bacteria isolated from animals. Approved standard, 3rd ed. Wayne: CLSI; 2008. CLSI Document M31-A3

Couñago, R.M., Davlieva, M., Strych, U., Hill, R.E. and Krause, K.L., 2009. Biochemical and structural characterization of alanine racemase from *Bacillus anthracis* (Ames). BMC structural biology 9, 53.

Dal Pozzo, M., Viégas, J., Santurio, D.F., Rossatto, L., Soares, I.H., Alves, S.H. and Costa, M.M.d., 2011. Atividade antimicrobiana de óleos essenciais de condimentos frente a Staphylococcus spp. Isolados de mastite caprina. Ciência Rural 11, 54–56.

Duque, E., Daddaoua, A., Cordero, B.F., De la Torre, J., Antonia Molina-Henares, M. and Ramos, J.L., 2017. Identification and elucidation of in vivo function of two alanine racemases from *Pseudomonas putida* KT2440. Environmental microbiology reports 9, 581–588.

Francois, J.A. and Kappock, T.J., 2007. Alanine racemase from the acidophile *Acetobacter aceti*. Protein expression and purification 51, 39–48.

Guo, S., Mo, Z., Wang, Z., Xu, J., Li, Y., Dan, X. and Li, A., 2018. Isolation and pathogenicity of *Streptococcus iniae* in offshore cage-cultured Trachinotus ovatus in China. Aquaculture 492, 247–252.

Hashimoto, A., Nishikawa, T., Oka, T., Takahashi, K. and Hayashi, T., 1992. Determination of free amino acid enantiomers in rat brain and serum by high-performance liquid chromatography after derivatization with N-tert.-butyloxycarbonyl-L-cysteine and o-phthaldialdehyde. Journal of Chromatography B: Biomedical Sciences and Applications 582, 41–48.

Higashi, T., 1958. Two types of hydroquinone oxidase of *Pseudomonas aeruginosa*. The Journal of Biochemistry 45, 785–793.

Hunter, R.C. and Newman, D.K., 2010. A putative ABC transporter, hatABCDE, is among molecular determinants of pyomelanin production in *Pseudomonas aeruginosa*. Journal of bacteriology 192, 5962–5971.

Im, H., Sharpe, M.L., Strych, U., Davlieva, M. and Krause, K.L., 2011. The crystal structure of alanine racemase from *Streptococcus pneumoniae*, a target for structure-based drug design. BMC microbiology 11, 116.

Ju, J., Xu, S., Wen, J., Li, G., Ohnishi, K., Xue, Y. and Ma, Y., 2009. Characterization of endogenous pyridoxal 5′-phosphate-dependent alanine racemase from *Bacillus pseudofirmus* OF4. Journal of bioscience and bioengineering 107, 225–229.

Kawakami, R., Ohshida, T., Sakuraba, H. and Ohshima, T., 2018. A Novel PLP-Dependent Alanine/Serine Racemase From the Hyperthermophilic *Archaeon Pyrococcus horikoshii* OT-3. Frontiers in microbiology 9, 1481.

Kim, M.G., Sbych, U., Krause, K., Benedik, M. and Kohn, H., 2003a. Evaluation of amino-substituted heterocyclic derivatives as alanine racemase inhibitors. Medicinal chemistry research 12, 130–138.

Kim, M.G., Strych, U., Krause, K., Benedik, M. and Kohn, H., 2003b. N (2)-Substituted D, L-Cycloserine Derivatives. The Journal of antibiotics 56, 160–168.

Lee, Y., Mootien, S., Shoen, C., Destefano, M., Cirillo, P., Asojo, O.A., Yeung, K.R., Ledizet, M., Cynamon, M.H. and Aristoff, P.A., 2013. Inhibition of mycobacterial alanine racemase activity and growth by thiadiazolidinones. Biochemical pharmacology 86, 222–230.

Liu, D., Liu, X., Zhang, L., Jiao, H., Ju, J. and Zhao, B., 2015. Biochemical characteristics of an alanine racemase from *Aeromonas hydrophil* HBNUAh01. Microbiology 84, 202–209.

Liu, D., Zhang, T., Wang, Y., Muhammad, M., Xue, W., Ju, J. and Zhao, B., 2019. Knockout of alanine racemase gene attenuates the pathogenicity of *Aeromonas hydrophila*. BMC microbiology 19, 72.

Liu, S., Wei, Y., Zhou, X., Zhang, K., Peng, X., Ren, B., Chen, V., Cheng, L. and Li, M., 2018. Function of alanine racemase in the physiological activity and cariogenicity of *Streptococcus mutans*. Scientific reports 8, 5984.

Muhammad, M., Zhang, T., Gong, S., Bai, J., Ju, J., Zhao, B., Liu, D., 2019. *Streptococcus iniae*: A Growing Threat and Causative Agent of Disease Outbreak in Farmed Chinese Sturgeon (*Acipenser sinensis*). Pakistan Journal of Zoology Accepted manuscript.

Nachbauer, C.A., James, J.H., Edwards, L.L., Ghory, M.J. and Fischer, J.E., 1984. Infusion of branched chain-enriched amino acid solutions in sepsis. The American journal of surgery 147, 743–752.

Nakatani, Y., Opel-Reading, H.K., Merker, M., Machado, D., Andres, S., Kumar, S.S., Moradigaravand, D., Coll, F., Perdigão, J. and Portugal, I., 2017. Role of alanine racemase mutations in *Mycobacterium tuberculosis* D-cycloserine resistance. Antimicrobial agents and chemotherapy 61, e01575–17.

Palumbo, E., Favier, C.F., Deghorain, M., Cocconcelli, P.S., Grangette, C., Mercenier, A., Vaughan, E.E. and Hols, P., 2004. Knockout of the alanine racemase gene in *Lactobacillus plantarum* results in septation defects and cell wall perforation. FEMS microbiology letters 233, 131–138.

Patrick, W.M., Weisner, J. and Blackburn, J.M., 2002. Site-directed mutagenesis of Tyr354 in *Geobacillus stearothermophilus* alanine racemase identifies a role in controlling substrate specificity and a possible role in the evolution of antibiotic resistance. Chembiochem 3, 789–792.

Qiu, W., Zheng, X., Wei, Y., Zhou, X., Zhang, K., Wang, S., Cheng, L., Li, Y., Ren, B. and Xu, X., 2016. d-Alanine metabolism is essential for growth and biofilm formation of *Streptococcus mutans*. Molecular oral microbiology 31, 435–444.

Ray, S., Das, S., Panda, P.K. and Suar, M., 2018. Identification of a new alanine racemase in *Salmonella Enteritidis* and its contribution to pathogenesis. Gut pathogens 10, 30.

Saavedra, M.J., Guedes-Novais, S., Alves, A., Rema, P., Tacão, M., Correia, A. and Martínez-Murcia, A., 2004. Resistance to β-lactam antibiotics in *Aeromonas hydrophila* isolated from rainbow trout (Onchorhynchus mykiss). International Microbiology 7, 207–211.

Scaletti, E.R., Luckner, S.R. and Krause, K.L., 2012. Structural features and kinetic characterization of alanine racemase from *Staphylococcus aureus* (Mu50). Acta Crystallographica Section D: Biological Crystallography 68, 82–92.

Seow, T.K., Inagaki, K., Tamura, T., Soda, K. and Tanaka, H., 1998. Alanine racemase from an *acidophile, Acidiphilium organovorum*: purification and characterization. Bioscience, biotechnology, and biochemistry 62, 242–247.

Shrestha, R., Lockless, S.W. and Sorg, J.A., 2017. A *Clostridium difficile* alanine racemase affects spore germination and accommodates serine as a substrate. Journal of Biological Chemistry 292, 10735–10742.

Soda, K. and Tanizawa, K., 1990. Thermostable Alanine Racemase. Annals of the New York Academy of Sciences 585, 386–393.

Strych, U., Davlieva, M., Longtin, J.P., Murphy, E.L., Im, H., Benedik, M.J. and Krause, K.L., 2007. Purification and preliminary crystallization of alanine racemase from *Streptococcus pneumoniae*. BMC microbiology 7, 40.

Tassoni, R., van der Aart, L.T., Ubbink, M., Van Wezel, G.P. and Pannu, N.S., 2017. Structural and functional characterization of the alanine racemase from *Streptomyces coelicolor* A3 (2). Biochemical and biophysical research communications 483, 122–128.

Tavares, G.C., de Queiroz, G.A., Assis, G.B.N., Leibowitz, M.P., Teixeira, J.P., Figueiredo, H.C.P. and Leal, C.A.G., 2018. Disease outbreaks in farmed Amazon catfish (*Leiarius marmoratus x Pseudoplatystoma corruscans*) caused by *Streptococcus agalactiae, S. iniae, and S. dysgalactiae*. Aquaculture.

Teulé, F., Cooper, A.R., Furin, W.A., Bittencourt, D., Rech, E.L., Brooks, A. and Lewis, R.V., 2009. A protocol for the production of recombinant spider silk-like proteins for artificial fiber spinning. Nature protocols 4, 341.

Toney, M.D., 2005. Reaction specificity in pyridoxal phosphate enzymes. Archives of biochemistry and biophysics 433, 279–287.

Wang, Y., Yang, C., Xue, W., Zhang, T., Liu, X., Ju, J., Zhao, B. and Liu, D., 2017. Selection and characterization of alanine racemase inhibitors against *Aeromonas hydrophila*. BMC microbiology 17, 122.

Wei, Y., Qiu, W., Zhou, X.-D., Zheng, X., Zhang, K.-K., Wang, S.-D., Li, Y.-Q., Cheng, L., Li, J.-Y. and Xu, X., 2016. Alanine racemase is essential for the growth and interspecies competitiveness of *Streptococcus mutans*. International journal of oral science 8, 231.

Yokoigawa, K., Kawai, H., Endo, K., Lim, Y.H., Esaki, N. and Soda, K., 1993. Thermolabile alanine racemase from a psychrotroph, Pseudomonas fluorescens: purification and properties. Bioscience, biotechnology, and biochemistry 57, 93–97.

